# Dorsal anterior cingulate cortex intrinsic functional connectivity linked to electrocortical measures of error-monitoring

**DOI:** 10.1101/2020.10.21.348649

**Authors:** Hayley Gilbertson, Lin Fang, Jeremy A. Andrzejewski, Joshua M. Carlson

## Abstract

The error-related negativity (ERN) is a response-locked event-related potential, occurring approximately 50 ms following an erroneous response at frontocentral electrode sites. Source localization and functional magnetic resonance imaging (fMRI) research indicate that the ERN is likely generated by activity in the dorsal anterior cingulate cortex (dACC). The dACC is thought to be a part of a broader network of brain regions that collectively comprise an error-monitoring network. However, little is known about how intrinsic connectivity within the dACC-based error-monitoring network contributes to variability in ERN amplitude. The purpose of this study was to assess the relationship between dACC functional connectivity and ERN amplitude. In a sample of 53 highly trait-anxious individuals, the ERN was elicited in a flanker task and functional connectivity was assessed in a 10-minute resting-state fMRI scan. Results suggest that the strength of dACC seeded functional connectivity with the supplementary motor area is correlated with the ΔERN (i.e., incorrect – correct responses) amplitude such that greater ΔERN amplitude was accompanied by greater functional coupling between these regions. In addition to the dACC, exploratory analyses found that functional connectivity in the caudate, cerebellum, and a number of regions in the error-monitoring network were linked to variability in ΔERN amplitude. In sum, ERN amplitude appears to be related to the strength of functional connectivity between error-monitoring and motor control regions of the brain.

## 1. Introduction

Nobody likes to make mistakes. Indeed, error commission is generally considered aversive and, when possible, avoided. Mistakes can be embarrassing and potentially harmful to one’s social status as well as potentially harmful to one’s physical health (Hajcak & Foti, 2008; Anna Weinberg, Riesel, & Hajcak, 2012). An aversion to errors is adaptive and crucial for survival—avoiding mistakes leads to more productive goal-relevant behavior. The detection of an error can lead to the modification or adjustment of behavior (e.g., increase in attentional resources) to optimize goal-related performance and decrease one’s chances of making future errors (Larson & Clayson, 2011; Taylor, Stern, & Gehring, 2007). On the other hand, past research and meta-analysis shows that heightened error-monitoring is related to anxious symptoms (Hajcak, McDonald, & Simons, 2003; Moser, Moran, Schroder, Donnellan, & Yeung, 2013; Weinberg, Kotov, & Proudfit, 2015), which indicates that it can be an important risk factor for the development and maintenance of anxiety (Meyer, Hajcak, Torpey-Newman, Kujawa, & Klein, 2015). Given the essential role of error-monitoring in both adaptive and maladaptive behavioral regulation, there has been great interest in its measurement and underlying neural mechanisms.

One of the most common ways to measure error-monitoring is through the use of an event related potential (ERP) known as the error-related negativity (ERN). The ERN is a response-locked ERP thought to reflect activation of an error detection system in the brain (Falkenstein, Hoormann, Christ, & Hohnsbein, 2000; Gehring, Goss, Coles, Meyer, & Donchin, 1993). The ERN has a negative amplitude occurring approximately 50ms after the commission of an erroneous response. ERN amplitude is proportionally linked to the significance of an error (Gehring et al., 1993; Hajcak, Moser, Yeung, & Simons, 2005). For example, ERN amplitudes are enhanced when task instructions prioritize accuracy over speed (Gehring et al., 1993). In addition, individual differences in sensitivity to errors are associated with ERN amplitude. For instance, individuals with heightened levels of anxiety (Hajcak, McDonald, & Simons, 2003; Moser, Moran, Schroder, Donnellan, & Yeung, 2013; Weinberg, Kotov, & Proudfit, 2015) and obsessive compulsive traits (Gehring, Himle, & Nisenson, 2000; Hajcak & Simons, 2002; Zambrano-Vazquez & Allen, 2014) exhibit larger ERN amplitudes. Accordingly, the ERN appears to be to reflect an error detection system sensitive to error intensity (Hajcak et al., 2005). While the ERN has been suggested to be functionally linked to error-monitoring, the ERN has also been associated with other theoretical frameworks, such as conflict monitoring theory (where errors are a specific type of conflict between actual and desired outcomes; see Larson, Clayson, & Clawson, 2014 for a review), reinforcement learning theory (Holroyd & Coles, 2002), mismatch theory, and motivational significance theory (see Olvet & Hajcak 2008 for a review). Additionally, some research suggests that the association between enhanced ERN amplitudes and anxiety reflects adaptive cognitive control rather than error-monitoring (Moser et al., 2013).

Research regarding the underlying neural mechanisms of error-monitoring has shown that the more caudal (or dorsal) region of the anterior cingulate cortex (dACC) plays a role in conflict, error, and action monitoring, while integrating emotional and motivational factors to maintain or adjust behavior (Taylor et al., 2007). For example, in functional magnetic resonance imaging (fMRI) studies of error-monitoring, the dACC displays increased activity immediately following incorrect, relative to correct, responses (Carter et al., 1998; Taylor et al., 2007; van Veen & Carter, 2002). Consistent with fMRI research, early source localization studies have identified this same dACC region and surrounding areas of the posterior medial prefrontal cortex (mPFC) as the likely generator of the ERN (Dehaene, Posner, & Tucker, 1994). Subsequent source localization research has largely confirmed this initial estimation by consistently localizing the ERN to the dACC (Herrmann, Römmler, Ehlis, Heidrich, & Fallgatter, 2004; O’Connell et al., 2007; Van Boxtel, Van Der Molen, & Jennings, 2005). Moreover, combined EEG-fMRI studies, provide multimodal evidence that ERN amplitude is linked to dACC activity at both the subject and single trial levels (Debener et al., 2005; Edwards, Calhoun, & Kiehl, 2012; Grutzmann et al., 2016; Iannaccone et al., 2015). Taken together, previous neuroimaging research provides strong convergent evidence that the ERN originates from activity in the dACC as part of an error-monitoring system.

In addition to the dACC, fMRI-based studies have identified increased error-related activity in a number of other cortical regions including the supplementary motor area (SMA), dorsolateral PFC (dlPFC), ventrolateral PFC (vlPFC), anterior insula (AI), inferior parietal lobule (IPL), and ventral/rostral ACC (Taylor et al., 2007) as well as subcortical brain structures such as the amygdala (Polli et al., 2008; Pourtois et al., 2010). These regions are collectively thought to form an error-monitoring network (Bastin et al., 2016; Taylor et al., 2007), in which the dACC can be seen as a central hub. The dACC receives reward vs. punishment valence-related information from the amygdala and vACC (Rolls, 2019), and mediates action-outcome monitoring and learning by projecting to the SMA and other motor regions, where motor outputs are regulated (Nachev, Kennard, & Husain, 2008; Rolls, 2019). The dACC also monitors and sends conflict information to lPFC, where increased cognitive control can be exerted to improve task performance (Cavanagh, Meyer, & Hajcak, 2017; Taylor et al., 2007). In addition, other regions within the network, such as the AI, together with the dACC are involved in the appraisal and regulation of emotional salience (Seeley et al., 2007). In sum, the dACC is a critical structure within a broader network of brain structures involved in error-monitoring and behavioral adjustment.

Yet, little is known about the strength of intrinsic connectivity between this error-monitoring network and electrophysiological measures of error-monitoring—in particular the ERN. Resting state functional connectivity (rsFC) measures the temporal reliance of activation between brain areas measured during a resting (i.e., non-task) state (Deco, Jirsa, & McIntosh, 2011). One common approach for assessing rsFC identifies the neural time course in a seed region and assesses concurrent/correlated neural activation in distributed brain regions. Identifying brain regions that are intrinsically connected could provide evidence of a network associated with an inherent cognitive process (Deco et al., 2011). To the best of our knowledge, no research has explored the relationship between rsFC and ERN amplitude to understand how intrinsic connectivity relates to error-monitoring.

To address this knowledge gap, this study assessed the relationship between ERN amplitude and rsFC seeded in the dACC. Given that individuals with heightened levels of anxiety are more sensitive to errors and tend to have larger ERNs (Moser et al., 2013), we used a sample of participants selected for high levels of trait anxiety. That is, we aimed to initially test the association between variability in ERN amplitudes and dACC seeded rsFC in a sample of individuals known to robustly produce ERN amplitudes, which would lead to greater ERN variability. Our primary hypothesis was that increased ERN (i.e., incorrect – correct, ΔERN) amplitudes would correlate with increased functional connectivity between the dACC (seed region) and distributed brain regions. In addition, exploratory analyses were performed using secondary seeds within the error-monitoring network (i.e., SMA, LPFC, AI, amygdala, and IPS), as well as brain regions that are not part of the error-monitoring network, but are also involved in error-related aspects of motor control and thus might indirectly contribute to the generation of ERN, such as the cerebellum and caudate (Mathalon, Whitfield, & Ford, 2003; Menon, Adleman, White, Glover, & Reiss, 2001).

## 2. Methods

### 2.1 Participants

Fifty-three individuals (male = 20, *M* = 22.74, *SD* = 5.71) were recruited from the university and surrounding community (18 – 38 years of age). Prior to the study, participants provided written informed consent. Participants included in this report were recruited for a clinical trial assessing the effects of attention bias modification on changes in brain structure (NCT03092609). Participants and data included here are from a subsample of participants from the pre-training session. We ran sensitivity analysis using G*Power (version 3.1.9.2) with α = .001, power = .80, total sample size = 53, which indicate that our study was powered to detect effect sizes of ρ ≥ 0.50.

### 2.2 Screening Procedure

To be included in the study, participants were screened to meet the following criteria (1) right-handed, (2) 18 – 42 years of age, (3) normal (or corrected to normal) vision, (4) no current psychological treatment, (5) no recent history of head injury or loss of consciousness, (6) no current psychoactive medications, (7) not claustrophobic, (8) not pregnant, (9) no metal in the body or other MRI contraindications, (10) trait anxiety scores ≥ 40 on the STAI-T (see below for questionnaire information), and (11) attentional bias scores ≥ 7ms in the dot-probe task^1^. If all inclusion criteria were met, participants were scheduled for EEG and fMRI sessions.

### 2.3 State-Trait Anxiety Inventory

State and trait anxiety were measured with the Spielberger State-Trait Anxiety Inventory (STAI; Spielberger, Gorsuch, & Lushene, 1970). The STAI has two subscales: the STAI-S assess how anxious one currently feels (i.e., state anxiety) and STAI-T assess how anxious one generally feels (i.e., trait anxiety). Each subscale has 20 items and participants were asked to rate their answer on a 4-point Likert scale. The total scores for each subscale range from 20-80. A score of 40 or higher has been linked to clinically relevant levels of anxiety in adults (> 17 years of age) and was a baseline threshold for inclusion in our study (Emons, Habibović, & Pederson, 2019; Julian, 2011; Spielberger et al., 1970).

### 2.4 Flanker Task

The Flanker Task is the most common and effective measure used to elicit an ERN. We used a modified flanker task programmed in E-Prime 3.0 (Psychology Software Tools, Sharpsburg, PA) presentation software. Participants were seated 59 cm from the computer screen. Responses were recorded with a Chronos response box (Psychology Software Tools, Sharpsburg, PA) placed in front of the computer monitor. As displayed in Figure 1, each trial began with a central fixation point (i.e., a white plus sign in the middle of a black screen) for 1000ms. Following the fixation, the screen displayed five white arrows centered on a black background for 200ms. Participants were instructed to indicate the direction of the center arrow (left or right) using the corresponding button on a Chronos response box (i.e., first button for left and second button for right). Half of the trials were compatible (i.e., < < < < < or > > > > >) and half incompatible (i.e., < < > < < or > > < > >). Each trial was followed by an inter-trial interval of 1000 to 1400 ms. The task included one practice block (20 trials) and seven experimental blocks. Each experimental block included 60 randomly presented trials. In total there were 420 trials (210 compatible and 210 incompatible). To maximize the validity of the ERN, participants were required to sustain an accuracy level of 75% to 90% for each block (Larson, Baldwin, Good, & Fair, 2010; Olvet & Hajcak, 2009; Pontifex et al., 2010).

**Fig. 1.**
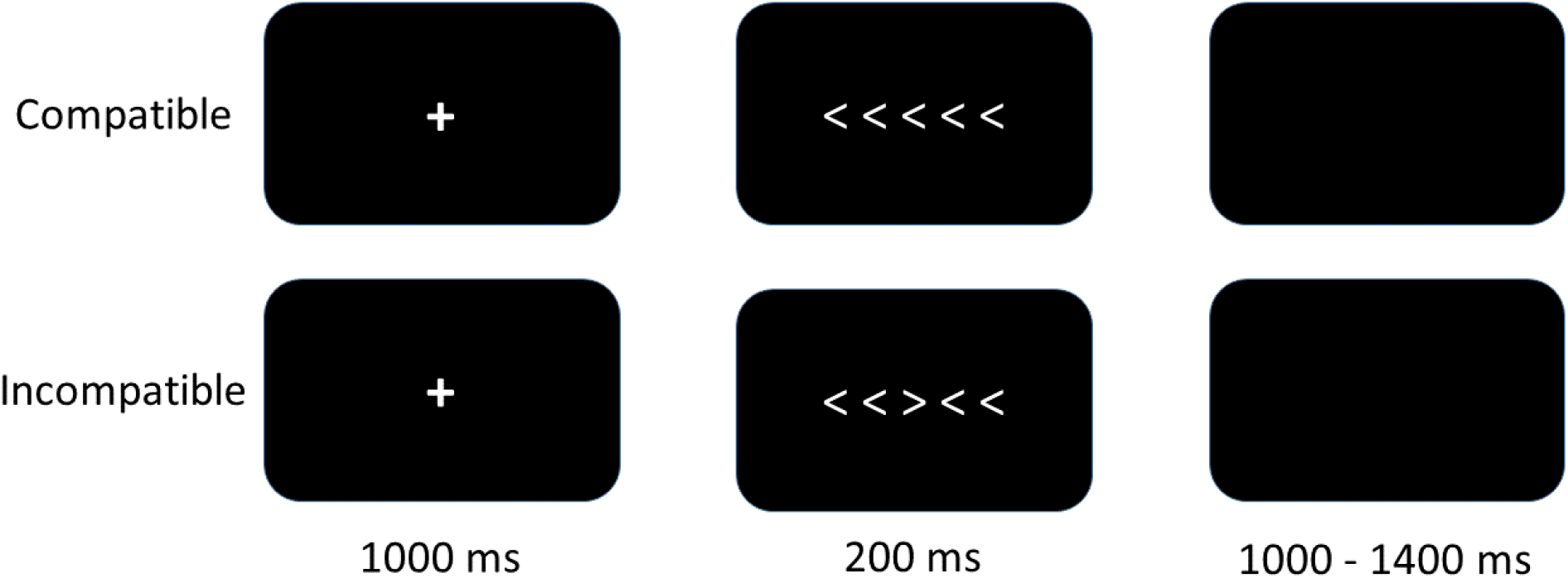
Example of the flanker task. During compatible trials, the center arrow and the peripheral/flanker arrows point to the same direction. In the incompatible trials, the center arrow points to the opposite direction of the peripheral/flanker arrows.

### 2.5 Neuroimaging Data-Acquisition and Analysis

#### 2.5.1 EEG Acquisition

The EEG data was recorded using a 64-channel EGI system (Electrical Geodesics Inc., Eugene, OR) with a 500 Hz sampling rate and reference at the vertex (Cz). Similar to previous research using the EGI system, electrode impedance levels were kept below 75 kΩ (Rizer, Aday, & Carlson, 2018; Tunison, Sylvain, Sterr, Hiley, & Carlson, 2019). Any channels with an impedance level over 75 and/or displaying other issues during the EEG session were logged.

#### 2.5.2 fMRI Acquisition

Functional MRI data were collected through a 1.5 Tesla General Electric whole-body scanner. Participants underwent a 10-minute resting state scan; during which they were instructed to relax and remain awake for the entirety of the scan. 240 functional volumes were collected using a T2* weighted gradient echo pulse sequence (TR = 2500 ms, TE = 35 ms, flip angle = 90°, FOV = 220, matrix = 64 × 64, voxel size = 3.4 mm × 3.4 mm, slice thickness = 5 mm). To co-register functional and structural images, high-resolution 3D FSPGR T1-weighted images were also obtained (TR/TI = min/450, flip angle = 9°, FOV = 250, matrix =256 × 256, voxel size = 0.98 × 0.98 mm, slice thickness = 1.2 mm).

#### 2.5.3 EEG Data Analysis

EEG data were preprocessed using EEGLAB toolbox v2019.0 (Delorme & Makeig, 2004) and ERPLAB toolbox V7.0.0 (Lopez-Calderon & Luck, 2014). EEG data were re-referenced using the averaged mastoid electrodes, and bandpass filtered (30 Hz low-pass, 0.1 Hz high-pass). Data were segmented from −500ms to 500ms around the participants’ response and split into correct versus incorrect categories. Data were baseline corrected (−400 ms to −200 ms). An independent component analysis (ICA) was then performed on segmented EEG data. After visual inspection of the scalp distributions and activity power spectrum of components for each participant, the components with obvious eyeblink and muscle artifacts were removed (Beatty, Buzzell, Roberts, & McDonald, 2020). In further artifact rejection, a step function was used to detect voltage exceeding 100 μV within the time period between 500 ms before and 200 ms after the response (Carr, Fitzroy, Tierney, White-Schwoch, Kraus, 2017; Schönhammer, Grubert, Kerzel, & Becker, 2016). For an individual’s average ERN to be valid, eight artifact-free incorrect response segments were required (Olvet & Hajcak, 2009; Pontifex et al., 2010). After the ICA and bad channel replacement operations, all the participants met the criteria and were thus included in further analysis so that their segment averages were combined into a grand average. Consistent with previous ERN research (e.g., Olvet & Hajcak, 2009; Pontifex et al., 2010), the mean amplitude of the frontocentral electrode, FCz, was extracted 0-100 ms post-response for each participant.

#### 2.5.4 fMRI Data Analysis

To convert DICOM files to NIfTI files, the fMRI scans were first imported through MRIcroGL (https://www.nitrc.org/projects/mricrogl), whereas the structural scans were imported using Statistical Parametric Mapping (SPM) 12 toolbox (Penny, Friston, Ashburner, Kiebel, & Nichols, 2011). The NifTI files were then preprocessed through the functional connectivity (CONN) toolbox (http://www.nitrc.org/projects/conn) in MATLAB (Math Works, Natick, MA). Images were realigned to correct for any head movement and then resliced to match the timing of the first image. In order to control for the effects of head motion, subject motion was calculated and removed automatically in the artifact detection procedure (subject-motion threshold = 0.9 mm, global-signal z-value threshold = 5). All the participants had more than 50% scans remaining; therefore all participants were included in further analysis. The six motion parameters and their corresponding first-order derives were taken into account in the regression models using valid scans. After being segmented into gray matter, white matter, and cerebrospinal fluid (CSF), images were normalized to the Montreal Neurological Institute (MNI) space and smoothed with an 8-mm Gaussian kernel of full width at half-maximum. Seed-based analyses were performed by calculating the Pearson’s correlation coefficients that represent the correlation between the time course of each seed area and all the voxels across the brain for each participant at the first-level of general linear model analysis. The correlation coefficients were then Fisher transformed into Z score, which was used in the second-level analysis with subject and ΔERN entered as covariates. Our primary analyses used the dACC (salience network: *xyz* = 0, 22, 35) seed in CONN and secondary analyses utilized the following bilateral seeds: SMA, LPFC, AI, amygdala, IPS, cerebellum, and caudate. All the ROIs have been derived using CONN’s default ROIs, which are defined from Harvard-Oxford Atlas (Desikan et al., 2006; Frazier et al., 2005; Makris et al., 2006) and Automated Anatomical Labeling Atlas (Tzourio-Mazover et al., 2002). All the results were initially thresholded at *p* < .001 (uncorrected) with a minimum cluster size of 20 voxels. Our primary analysis used a family wise error rate (FWE) cluster level corrected threshold of *p* < .05. Secondary (exploratory) analyses used the more liberal (uncorrected) threshold of *p* < .001.

## 3. Results

### 3.1 Behavior

The error rates of the Flanker task were *M* = 12.20%, *SD* = 4.88%, ranging from 4% to 32%. The reaction time (in milliseconds) of the correct trials (*M* = 375 ms, *SD* = 52) were significantly greater than the incorrect trials (*M* = 316 ms, *SD* = 58), *t* (52) = 18.81, *p* < .001. The average trait anxiety level was 53.25 (*SD* = 7.84), ranging from 40 to 70.

### 3.2 ERN

As displayed in Figure 2, ERN amplitudes were larger for incorrect compared to correct trials during the flanker task, *t*(52) = 6.97, *p* < .001; ΔERN (μV): *M* = −4.96, *SD* = 5.18. The relationship between ΔERN amplitude and trait anxiety was not significant, *r* = .24, *p* = .09.

**Fig. 2.**
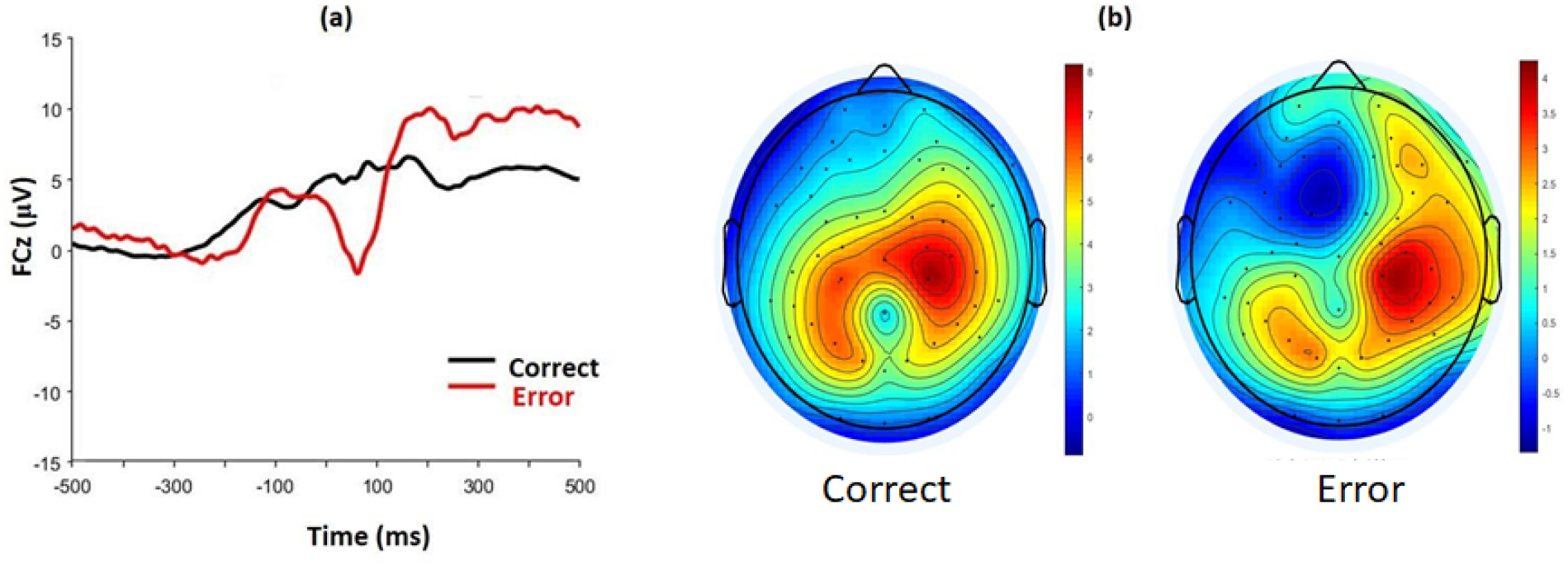
(a). Grand average waveforms for correct and incorrect response at electrode FCz for the flanker task. (b). Scalp topography for correct and incorrect response between 0-100 ms post-response.

### 3.3 ERN - rsFC

As displayed in Figure 3, the dACC seed resulted in a significant negative association (*r* = −.52) between ΔERN amplitude and the strength of rsFC between the dACC and the bilateral SMA, *xyz* = 10, −12, 70, *k* = 262, *z* = 3.63, *p*_*cluster FWE*_ < .05. Greater ΔERN amplitude (i.e., more negative) was associated with greater connectivity between the dACC and SMA. The extracted dACC-SMA connectivity was also associated with ERN amplitude (*r* = −.38, *p* < .01), residual ERN (*r* = −.50, *p* < .001), but not with CRN (*r* = .17, *p* = .23). The association between ΔERN amplitude and dACC-SMA connectivity was not significantly different between females (*r* = - .41, *p* < .05) and males (*r* = −.63, *p* < .01), *z* = 1.01, *p* = .31. After taking age, accuracy rate, and trait anxiety level into account, this association was still robust, *r* = −.54, *p* < .001. No other dACC-seeded connections were related to ΔERN amplitude at the FWE cluster level. The results of exploratory analyses performed using secondary seeds can be found in the Supplementary Table.

**Fig. 3.**
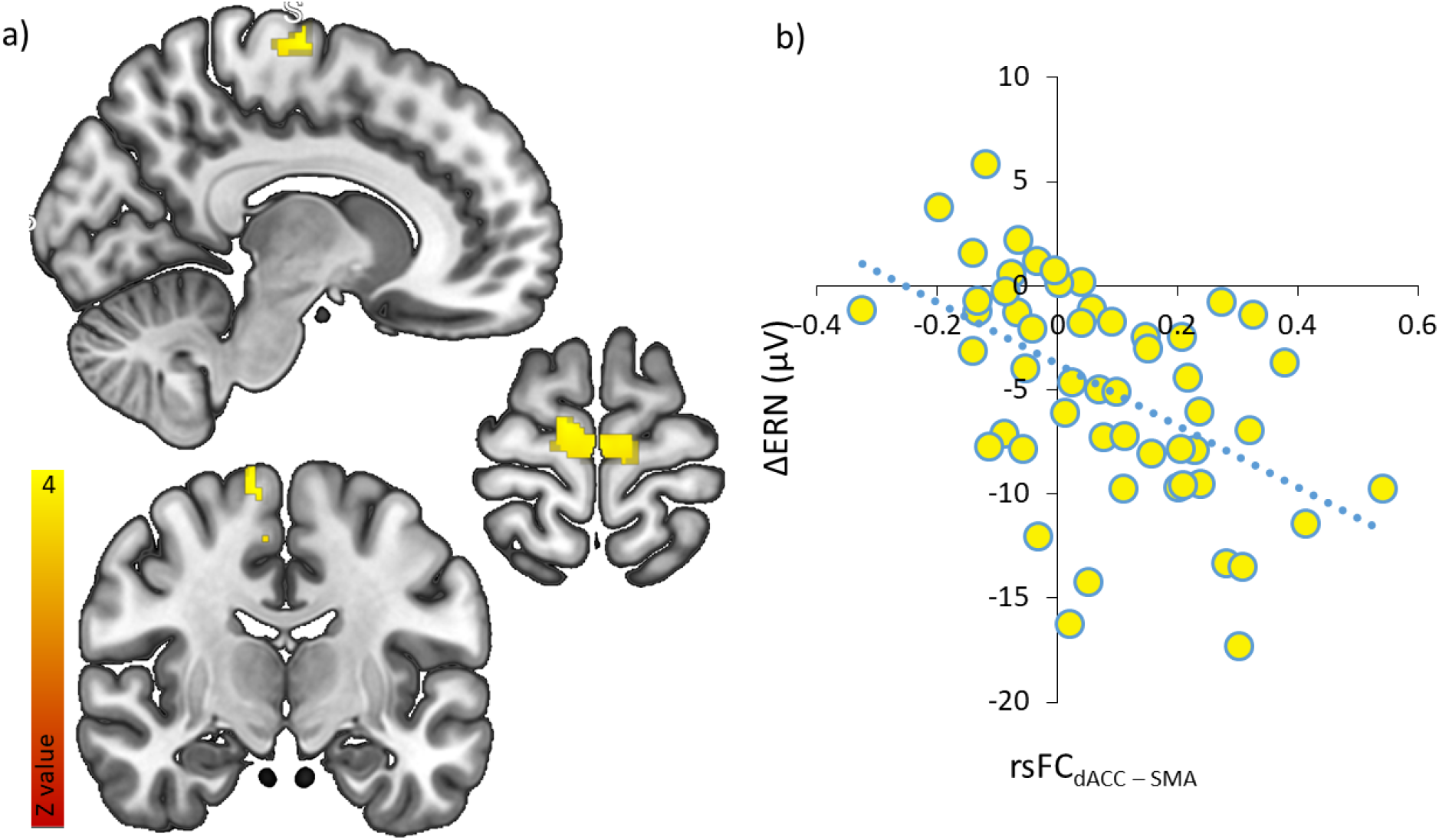
(a). dACC (0, 22, 35) seeded rsFC correlated with increased ΔERN amplitude in the bilateral motor cortex. (b). Scatterplot displaying the functional connectivity between the dACC and SMA (10, −12, 70) and its correlation with ΔERN.

## 4. Discussion

In a sample of highly anxious individuals we measured electrocortical measures of error-monitoring (i.e., the error-related negativity; ERN) during a flanker task. We additionally collected measures of resting-state functional connectivity (rsFC) within a dACC based “error-monitoring network”. We aimed to assess the relationship between ΔERN amplitude and intrinsic connectivity. Our flanker task was effective in eliciting the ERN as amplitudes were larger for incorrect, relative to correct, responses (see Figure 2). We present novel evidence that ΔERN amplitude is related to intrinsic connectivity between the dACC and SMA—highlighting an important link between error-monitoring and motor control regions in contributing to variability in ERN amplitude (see Figure 3). Beyond the dACC, exploratory analyses identified that rsFC between other brain regions was also related to ΔERN amplitude. For example, rsFC between the cerebellum and thalamus, as well as the caudate and precuneus correlated with ΔERN amplitude suggesting that variability in ERN amplitude is related to connectivity in a broader network of brain regions (see Supplementary Table). In short, we present the first evidence (that we are aware of) demonstrating that variability in ΔERN is related to rsFC in a dACC based error-monitoring network.

ΔERN amplitude was found to correlate with the strength of rsFC between the dACC and the SMA. In particular, greater rsFC between the dACC and SMA was linked to larger ΔERN amplitudes. This association was still robust, after taking into account age, accuracy rate, and trait anxiety. In addition, greater rsFC between the dACC and SMA was correlated with (raw incorrect) ERN amplitudes as well as the residual ERN, but not (raw correct) CRN amplitudes. This pattern of results suggests that the link between ΔERN amplitudes and dACC seeded rsFC is primarily being driven by the neural activity on incorrect trials related to error-monitoring and error-detection. Although previous research has shown that the relationship between ERN and anxiety is moderated by sex such that this relationship is stronger in females (Moser, Moran, Kneip, Schroder, and Larson, 2016), we did not see significant differences between females and males in the strength of the correlation between rsFC and ERN amplitude. Thus, the relationship between ERN amplitude and the strength of rsFC between the dACC and SMA is robust to a number of potential confounds, methodological quantifications of the ERN, and participant sex.

Given that the ERN is a response-locked ERP, the association between the dACC and SMA is not surprising, although previously unknown. Indeed, the dACC has been implicated in integrating valence-related (i.e., an aversive error/mistake) and location-related (i.e., the spatial location of the preceding movement) information to mediate action-outcome learning and behavior (Rolls, 2019). Specifically, within the framework of detection and correction of an erroneous response or action, the dACC is thought to monitor behavior in relation to an individual’s current behavioral objectives (Taylor et al., 2007), while the SMA is involved in the planning and execution of a behavioral response (Nachev et al., 2008). Our findings may indicate that processes related to behavioral adjustment/compensation following an erroneous response are reflected in ΔERN amplitudes. This connection would be adaptive and may allow for increased action monitoring and action adjustment following a mistake.

This interpretation would be consistent with conflict monitoring theory, which posits that errors (i.e., a type of conflict) are detected and initiate regulatory control processes intended to correct the error (Larson, Clayson, & Clawson, 2014; Yeung, Botvinick, & Cohen, 2004). Although these regulatory control processes are often thought to involve prefrontal cognitive control, motor control processes also act to minimize erroneous responses. It should be noted, however, that we did not design our study to test opposing theories related to the functional significance of the ERN and other ERN theories (e.g., reinforcement learning theory; Holroyd & Coles, 2002) may also explain the pattern of results observed here. Regardless of the functionality underlying this association, individual differences in ΔERN amplitude are linked to differences in the connectivity strength between the dACC and motor regions.

Resting state networks are defined by the coactivation of brain regions during non-task states. This coactivation is thought to reflect a shared functional role of the brain structures (Deco et al., 2011). In addition, coactivation, or functional connectivity, is thought to indicate a direct or indirect structural pathway between the structures within the functional network (Deco et al., 2011). Indeed, previous research has shown that rsFC measures are highly correlated with diffusion tensor-weighted structural connectivity measures (Carlson, Cha, & Mujica-Parodi, 2013). The dACC and SMA are anatomically connected and this connection is thought to subserve action-outcome behaviors (Rolls, 2019). Thus, rsFC between the dACC and SMA may reflect an underlying anatomical connection between these structures, which is related to ΔERN amplitude. However, future research will be needed to determine the extent to which the link between ΔERN amplitude and dACC-SMA connectivity is mediated by the strength of a direct anatomical connection between these structures.

Beyond dACC seeded rsFC, exploratory analyses suggest that variability in the intrinsic connectivity of other structures in the broader error-monitoring network are related to ERN amplitude (see Supplementary Table). For example, we found that greater ΔERN amplitude was associated with greater strength of connectivity between the lPFC and regions of the mPFC as well as the lPFC and regions of the temporal cortex. Elevated connectivity between these structures may suggest a potential association between self-relevance evaluation and error-monitoring (Gusnard, Akbudak, Shulman, & Raichle, 2001; Ochsner et al., 2004; Raichle et al., 2001; Ralchle & Snyder, 2007; Taylor et al., 2007). Moreover, we found that greater rsFC between the cerebellum (crus 2) and the thalamus as well as greater rsFC between the caudate and precuneus were related to smaller ΔERN amplitudes. These associations provide evidence that the cerebellum and caudate could indirectly engage in error-related processes linked to the ERN (Holroyd & Coles, 2002; Mathalon et al., 2003). Taken together, our exploratory results indicate that variability in ERN amplitude may be related to rsFC in a broader error-monitoring network as well as brain regions involved in error-related aspects of motor control. Yet, future research is needed to corroborate these exploratory findings and further examine the specific role of these associations in the ERN-based error-monitoring system.

Although the current investigation only measured error-monitoring in subclinical anxiety, our results may have clinical implications. Anxiety disorders, obsessive compulsive disorder (OCD), and schizophrenia are all prevalent mental illnesses that are associated with abnormal error-monitoring (Gehring et al., 2000; Grutzmann et al., 2016; Hajcak et al., 2003; Hajcak & Simons, 2002; Moser et al., 2013; Polli et al., 2008; Weinberg et al., 2015; Zambrano-Vazquez & Allen, 2014). Therefore, it is important to understand the neural mechanisms that contribute to error-monitoring and abnormal error-monitoring in psychopathology. Our results suggest that differences in ΔERN amplitude are related to underlying differences in intrinsic connectivity, which may provide a more mechanistic explanation for abnormal error-monitoring as indexed by ERN amplitude. That is, at the neural level, variability in ΔERN amplitude may be the product of distinct alterations in the connectivity between nodes in the error-monitoring network (e.g., dACC-SMA). These distinct neural alterations are likely reflective of distinct alterations in sub-processes related to error-monitoring. For example, dACC-SMA connectivity would more likely reflect behavioral adjustment/compensation; whereas lPFC connectivity would likely reflect cognitive control (i.e., increased executive attention). Thus, our findings may indicate that ERN amplitudes in high trait anxiety individuals reflect adaptive processes related to the regulation and/or compensation of erroneous actions (Moser et al., 2013). These results may open the door for clinical research to explore the extent to which disorders associated with abnormal ERN amplitude have underlying abnormalities in network connectivity.

This study has several limitations. First, functional connectivity was assessed during a non-task state and it is therefore unclear if the rsFC observed here is reflective of processing that occurs during error-monitoring behavior. However, as indicated above, rsFC networks are believed to reflect networks of common activity and functional significance (Deco et al., 2011). Second, it should be noted that we did not directly measure structural connectivity since the main goal here was to assess the relationship between variability in ΔERN amplitude and individual differences in intrinsic connectivity. Therefore, we cannot conclude that the relationship between ΔERN amplitude and rsFC is mediated by the strength of underlying anatomical connectivity between the nodes in the network, even though rsFC measures have been shown to correlate with structural connectivity measures (Carlson et al., 2013). Finally, we selected a sample of individuals with high levels of trait anxiety based on prior research indicating that these individuals robustly produce an ERN. Therefore, like all studies, the extent to which our results can be generalized is based on the characteristics of our sample. Future research is needed to determine if the findings observed here generalize to non-anxious individuals, other age ranges, as well as other clinically relevant traits linked to elevated error-monitoring (e.g., OCD). Limitations aside, we provide strong evidence that the strength of rsFC between the dACC and SMA contributes to variability in ΔERN amplitude.

In sum, to the best of our knowledge, we present the first evidence that variability in ΔERN amplitude is related to rsFC in a dACC based error-monitoring network. ΔERN amplitude correlates with intrinsic connectivity between the dACC and SMA, which suggests an important link between error-monitoring and motor control processes in contributing to variability in ΔERN amplitude. Variability in the intrinsic connectivity of other nodes in error-monitoring network regions such as the SMA, lPFC, mPFC, and IPL were also related to ΔERN amplitude. These associations may reflect increased cognitive control and executive attention following error commission. In short, our findings indicate that variability in ΔERN amplitude is linked to underlying differences in intrinsic connectivity, providing a more mechanistic explanation for the network properties that give rise to variability in ERN amplitude.

## Author Note

Research reported in this publication was supported by the National Institute of Mental Health of the National Institutes of Health under Award Number R15MH110951. The content is solely the responsibility of the authors and does not necessarily represent the official views of the National Institutes of Health. We would like to thank Rourke Sylvain, Taylor Susa, and all the other students in the Cognitive × Affective Behavior & Integrative Neuroscience (CABIN) Lab at Northern Michigan University as well as Rochelle Milano and Annette Gustafson at UPHS-Marquette for assisting in the collection of this data.

## Footnotes

1 Note that the dot-probe task attention bias cutoff score was used for the larger attention bias modification study to only select individuals most likely to benefit from such training.

