## Supplementary Material for "Dorsal anterior cingulate cortex intrinsic functional connectivity linked to electrocortical measures of error-monitoring"

The analysis of caudate-seeded rsFC showed that the FC between caudate and precuneus was positively correlated with ΔERN, *r* = .57 (*p_cluster_* FWE < .05). The cerebellum-seeded rsFC revealed that the FC between cerebellum crus 2 and thalamus was positively correlated with ΔERN, *r* = .58 (p_cluster_ FWE < .05). Additionally, exploratory analyses using a more liberal threshold for bilateral seeds in the SMA, LPFC, AI, amygdala, and IPS can be found in the Supplementary Table S1.

### Table S1. Brain regions showing significant ERN related seed-based FC (voxel-wise *p* < .001 uncorrected)

| Brain Region | MNI  coordinates | | | Cluster Size  (voxels) | Peak  Z-score |
| --- | --- | --- | --- | --- | --- |
|  | X | Y | Z |  |  |
| ***Anterior cingulate cortex*** |  |  |  |  |  |
| Precentral gyrus | 22 | -24 | 54 | 59 | 3.47 |
| Superior parietal gyrus | 26 | -58 | 60 | 38 | 3.73 |
| ***Amygdala Left*** |  |  |  |  |  |
| Angular Gyrus | -52 | -52 | 34 | 43 | 3.47 |
| Superior Frontal Gyrus | -22 | 16 | 44 | 40 | 3.74 |
| Inferior frontal gyrus | -50 | 32 | 20 | 22 | 3.48 |
| Inferior temporal gyrus | 50 | -52 | -26 | 22 | 3.57 |
| ***Anterior Insula Left*** |  |  |  |  |  |
| Inferior frontal gyrus | 50 | 36 | 0 | 75 | 3.69 |
| Lateral occipital cortex | -40 | -66 | 54 | 40 | 3.85 |
| Precuneus cortex | 0 | -72 | 44 | 32 | 3.58 |
| ***Caudate Left*** |  |  |  |  |  |
| Precuneus Cortex | 14 | -48 | 54 | 451 | 4.31 |
| ***Cerebellum Crus2 Right*** |  |  |  |  |  |
| Thalamus | -8 | -32 | 10 | 353 | 4.10 |
| ***Intraparietal sulcus Left*** |  |  |  |  |  |
| Anterior cingulate cortex | 10 | 40 | 18 | 77 | 3.78 |
| ***Intraparietal sulcus Right*** |  |  |  |  |  |
| Middle frontal gyrus | -30 | 48 | 26 | 99 | 3.57 |
| Parahippocampal gyrus | -20 | 4 | -28 | 39 | 4.37 |
| Precuneus cortex | 8 | -78 | 46 | 32 | 3.67 |
| Cerebellum | -4 | -46 | -28 | 32 | 3.93 |
| Posterior cingulate gyrus | 6 | -42 | 16 | 23 | 3.49 |
| Precuneus | -14 | -56 | 62 | 20 | 3.46 |
| ***Lateral PFC Left*** |  |  |  |  |  |
| Superior temporal gyrus | -52 | 10 | -2 | 163 | 4.41 |
| Middle temporal gyrus | -62 | -64 | -8 | 163 | 4.03 |
| Frontal medial cortex | 4 | 50 | -18 | 40 | 3.66 |
| Temporal pole | -34 | 8 | -48 | 33 | 3.63 |
| Middle temporal gyrus | 68 | -16 | -12 | 29 | 3.91 |
| Superior frontal gyrus | 14 | 60 | 4 | 24 | 3.51 |
| Occipital pole | 14 | -102 | 6 | 23 | 3.58 |
| Insular cortex | 38 | -8 | -16 | 23 | 4.03 |
| ***Lateral PFC Right*** |  |  |  |  |  |
| Frontal medial cortex | 4 | 48 | -22 | 145 | 4.19 |
| Superior frontal gyrus | 0 | 48 | 32 | 76 | 3.74 |
| Inferior occipital gyrus | 38 | -80 | -16 | 62 | 3.55 |
| Superior frontal gyrus | -16 | 22 | 54 | 35 | 3.85 |
| ***Supplementary Motor Area Right*** |  |  |  |  |  |
| Middle Frontal Gyrus | -18 | 18 | 38 | 184 | 4.01 |

*Note:* ERN, Error-related negativity; FC, functional connectivity; MNI, Montreal Neurological Institute
